# Multiplane and Spectrally-Resolved Single Molecule Localization Microscopy with Industrial Grade CMOS Cameras

**DOI:** 10.1101/186544

**Authors:** Hazen P. Babcock

## Abstract

In this work we explore the use of industrial grade CMOS cameras for single molecule localization microscopy (SMLM). We show that the performance of these cameras in single imaging plane SMLM applications is comparable to much more expensive scientific CMOS (sCMOS) cameras. We show that these cameras can be used in more demanding biplane, multiplane and spectrally resolved SMLM applications. The 10-40× reduction in camera cost makes it practical to build SMLM setups with 4 or more cameras. In addition we provide open-source software for simultaneously controlling multiple CMOS cameras and for the reduction of the movies that are acquired to super-resolution images.

## Introduction

Super resolution imaging by localizing single flourescent molecules is popular due to its comparative simplicity. However it is generally accepted that SMLM requires the use of high end scientific cameras in order to detect the signals from single fluorescent dye molecules. This signal is typically of order 10-1000 photons per pixel at the magnifications used in most SMLM microscopy setups. The ideal camera should then have a high quantum efficiency (QE) for optimal conversion of photons to photo-electons (e-). This signal is also Poisson distributed so the camera readout noise should be less than 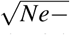 to minimally contribute to the total noise. In two of the three initial demonstrations of the SMLM approach^1–3^ and much of the subsequent work EMCCDs have been the detector of choice. These cameras have a maximum QE of over 80% and the EM gain stage amplifies the signal such that the relatively large (20e-to 40e-) readout noise of the CCD does not dominate the Poisson noise of the signal. More recently scientific CMOS (sCMOS) cameras have become popular due to their much higher readout rates, greater pixel counts and reduced cost^4-7^. Modern sCMOS cameras also have a maximum QE of over 80% and readout noises of ~1e-, which is neglible in comparison to the Poisson noise of the signal. Interestingly, high end industrial grade CMOS cameras are rapidly approaching the performance of sCMOS cameras. In particular it is easy to find industrial CMOS cameras with 65% maximum QE and read noises of ~2e-for $1.5k, an order of magnitude less than the approximately $20k cost of a typical sCMOS camera. This level of performance is sufficient for most types of SMLM imaging and there is little measurable difference in the quality of the final SMLM image from these cameras as compared to sCMOS cameras^8^.

This work follows earlier work that demonstrated several approaches to reducing the overall cost of a SMLM setup^8, 9^. However here we take advantage of the reduction in camera cost demonstrated in^8^ and use it to build a relatively inexpensive 4 camera setup. This is advantageous as it is simpler to construct a multiplane and/or multicolor imaging setup using a single camera per focal plane or color than it is to combine the focal planes or colors onto a single camera. In addition the final field of view of such a setup can be significantly larger as sharing the active area of a single camera is no longer necessary. Our setup can be quickly re-configured for different applications simply by swapping dichroics and/or beam splitters and adjusting lens positions. We demonstrate the use of this setup to acquire 2D SMLM images with a single camera, biplane and quadplane 3D SMLM images with 2 and 4 cameras respectively, and spectrally resolved SMLM (SR-STORM^10^) images with 4 cameras.

## Results

Fig. 1 is a diagram of the setup that was used in this work. It is built around an inverted microscope (TiU, Nikon) mounted on an optical table (RS2000, Newport). This microscope has an optical port selector that allows one to quickly change which output port the sample image is sent to. A single sCMOS camera (ORCA-Flash4.0, Hamamatsu Photonics) was mounted directly onto the left port of the microscope. An optical cage system was used to mount 4 industrial CMOS cameras (GS3-U3-51S5M-C, FLIR Imaging) onto the right port of the microscope. The setup can be configured to use from 1 to 4 of the CMOS cameras at once by adding or removing dichroic beam splitters in the fluorescent filter cube holders (**Sp** in Fig. 1, DFM1, Thorlabs). In order to adjust for the difference in pixel size between the two cameras (ORCA-Flash4.0 - 6.5 μm, GS3-U3-51S5M-C −3.45 μm) a 2× demagnifying lens pair was used (2*f* lens pair **L1**, **L2** in Fig. 1). A 60× 1.4NA oil immersion objective (CFI Plan Apo Lambda 60×, Nikon) was used for imaging, giving final pixel sizes of 108nm for the ORCA-Flash4.0 camera and 120 nm for the GS3-U3-51S5M-C cameras.

**Figure 1.**
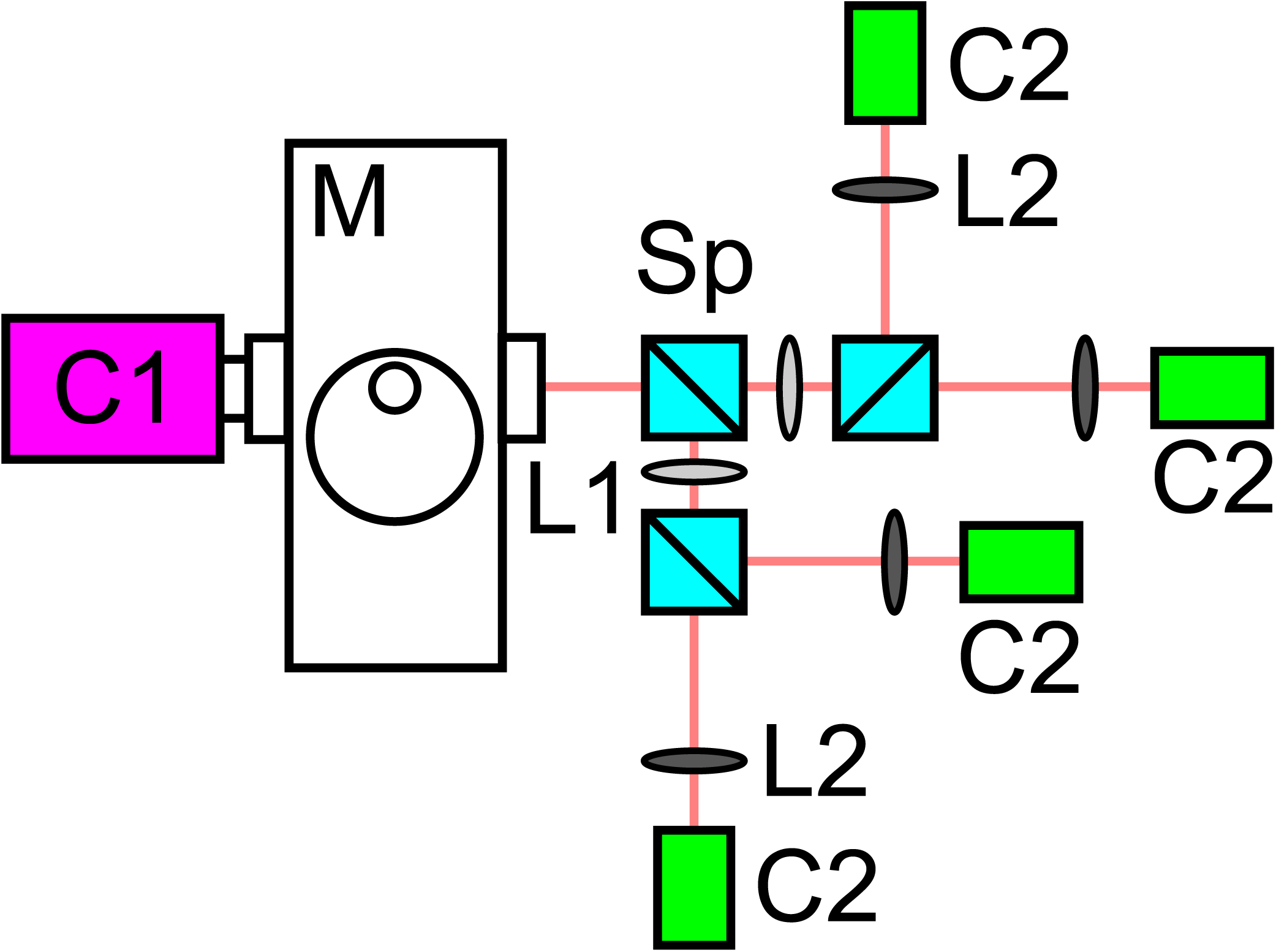
Schematic of the setup used for camera testing. **C1** (magenta) is a Hamamatsu ORCA-Flash4.0 camera, **M** is a Nikon TiU microscope, **Sp** (light blue) are flourescent filter cube holders, **L1** (light gray) *f* = 125 mm lenses, **L2** (dark gray) *f* = 60mm lenses and **C2** (green) are FLIR Imaging GS3-U3-51S5M-C cameras.

Other important components of the setup that are not shown in Fig. 1 include 1W 560nm and 1W 647 nm fiber lasers (2RU-VFL-P-1000-560-B1R, 2RU-VFL-P-1000-647-B1R, MPB photonics) used for fluorescence illumination. The lasers provided even illumination across a 40 μm diameter field of view, with typical powers for SMLM imaging of 3-6 kW/cm^2^ at the sample. A custom multi-band dichroic (zt405/488/561/647/752rpc, Chroma) and a custom multi-band emission filter (zet405/488/561/647-656/75, Chroma) were used for all of the experiments except SR-STORM. A 652 nm dichroic (FF652-DI01-25x36, AVRO Inc.) and a 647nm long pass emission filter (LP02-647RU-25, AVRO Inc.) were used for the SR-STORM experiments. 50/50 dichroic beam splitters (BSW10R, Thorlabs) were used to split the fluorescence emission between different CMOS cameras in the biplane and quadplane experiments. 670 nm, 695 nm and 720 nm dichroic filters (T670lpxr, T695lpxr and 720dcxr, Chroma) were used to spectrally separate the flourescence emission in the SR-STORM experiments. A 980 nm IR laser diode (LP980-SF15, Thorlabs), USB camera (DCC1645C, Thorlabs) and a piezo objective positioner (Nano-F100S, Mad City Labs) were used to build a IR reflectance based focus lock system that corrected for focal drift during data acquisition^11^.

The suitability of the CMOS camera for SMLM imaging was first evaluated by how well it could localize 0.1 μm 580/605 nm flourescent beads (F8801, Molecular Probes). This experiment was a side-by-side comparison of the relative performance of the CMOS and sCMOS cameras imaging exactly the same beads. Samples were prepared by sparsely and non-specifically immobilizing the beads onto a microscope coverslip. All of the fluorescence was imaged onto a single CMOS camera by removing the dichroics from the optical cage system. Pairs of 100 frame movies of the same field of view were taken at 100Hz with the sCMOS and CMOS cameras. The photon flux from the beads was adjusted to cover the range that is expected from single fluorescent dye molecules during SMLM imaging. The photo-bleaching of the beads during the acquisition of the two movies was neglible as these beads are very bright and require very little illumination laser power under these conditions. Analysis of the data was done by first finding the affine transform that best overlaid the images from the two different cameras. Then beads were identified and localized in movies from the sCMOS camera using a Python/C open-source implementation^12^ of the sCMOS analysis algorithm described in^5^. Next the bead fit locations were mapped to the CMOS camera and used as the starting points for fitting using the same sCMOS analysis algorithm. Beads were tracked through the movies by assigning all the beads that localized to within 2 pixel of a bead position in the first frame to the same track. Tracks where beads were missing in more than 10% of the frames in data from either of the cameras were discarded. Finally the average flux per bead in each frame was measured using aperture photometry. The median intensity of each frame was subtracted from the frame to minimize the contribution of background fluorescence. Then the integrated intensity in a 11×11 square pixel window centered on the track center was recorded and averaged across all of the frames.

Both cameras achieve the same localization precision as a function of bead intensity measured by the camera (Fig. 2A). This shows that the additional readout noise of the CMOS camera (1.6e-median) as compared to the sCMOS camera (0.93e-median) has little effect on the localization precision even at the lowest intensities tested here. The sCMOS camera however has a higher QE than the CMOS camera, so it will detect more photons per bead than the CMOS camera, and will demonstrate superior localization precision once this is corrected for. We measured the QE difference between the two cameras at the wavelength of the bead sample by plotting the bead intensity measured by the sCMOS camera versus that measured by the CMOS camera for each bead pair and fitting a line. We found that the QE of the CMOS camera was 67% of the QE of the sCMOS camera. Figure 2B is a plot of the localization precision with the intensity values from the CMOS camera corrected for this difference in QE. As expected, the localization precision of the sCMOS camera is about 25% better. It is also worth mentioning that this CMOS camera should perform as well as an EMCCD camera in SMLM applications with reasonably bright dyes. The excess noise added in the EMCCD gain stage is equivalent to reducing the signal by a factor of 2, so the performance of a perfect EMCCD would be approximately that of a CMOS camera with 50% QE^13^. Figure 2 also shows that the localization precision from both cameras, without correcting for the difference in QE, achieves the Cramer-Rao theoritical bound. The Cramer-Rao theoritical bound was calculated by numerical integration of equation 5 in^13^. A pixel size of 108 nm, a σ of 130 nm and background values determined as described in the preceding paragraph were used as the constants in this equation. The σ value was the average value returned by the sCMOS analysis algorithm for the width of the Gaussian fit to each localization. The apparent deviation at the highest intensities may be due to an underestimation of the image background.

**Figure 2.**
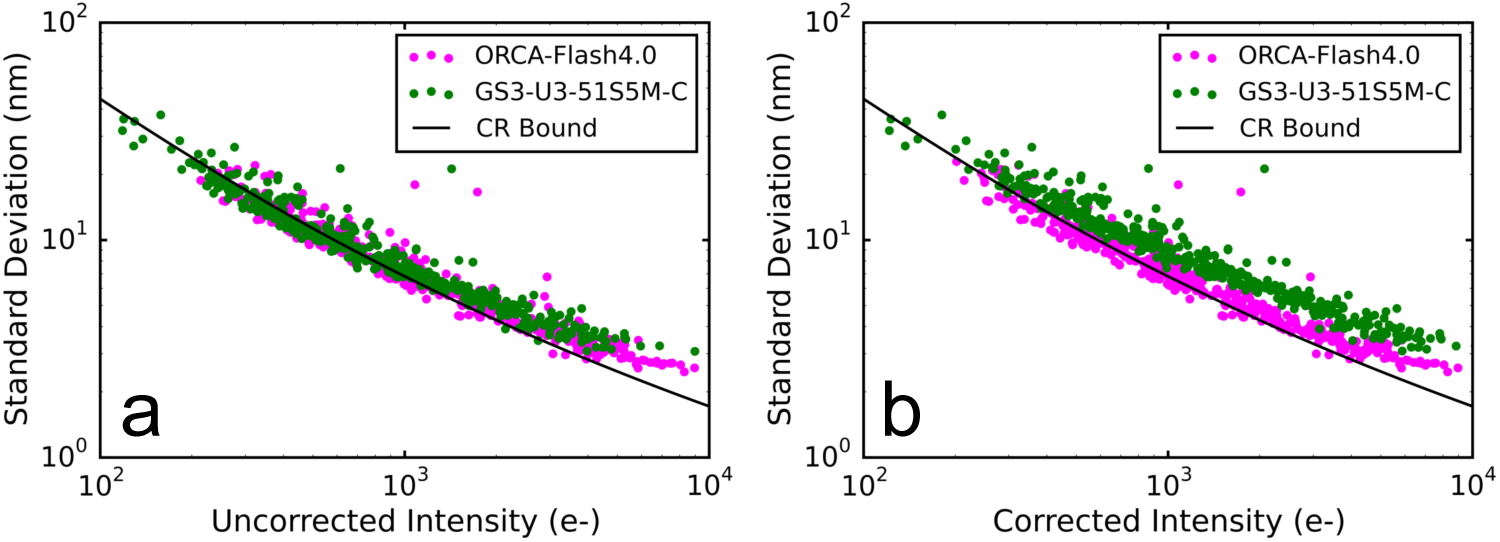
Comparison of localization precision measured using fluorescent beads (log-log plot). (**a**) Localization precision versus bead fluorescence intensity as measured by each camera.(**b**) Localization precision versus bead fluorescence intensity with the intensity reported by the GS3-U3-51S5M-C camera scaled by this camera’s sensitivity relative to the ORCA-Flash4.0 camera. The black line in both figures is the Cramer-Rao bound on the maximum localization precision.

The suitability of the CMOS camera for 2D SMLM imaging was next evaluated with a test sample consisting of microtubules labeled with the Alexa-647 fluorescent dye. In this experiment we took advantage of the fact that a single Alexa-647 dye molecule can be localized multiple times in order to acquire SMLM images of the same field of view with both the sCMOS and the CMOS cameras. In order to minimize the relative effects of photo-bleaching on the the final SMLM image we alternated imaging between the two cameras, acquiring relatively short movies with each camera before switching to the other camera. The procedure that was followed was an initial turn off phase followed by a 5k frame SMLM movie with the sCMOS camera. Then alternating 10k frame SMLM movies were taken with the CMOS camera and the sCMOS camera, and finally a 5k frame SMLM movie was taken with the sCMOS camera. The total SMLM movie length for each camera was 70k frames. All movies were taken at a frame rate of 100Hz with HILO illumination^14^. These movies were reduced to SMLM images with the sCMOS analysis algorithm mentioned above. A comparable number of localizations were identified with each camera, 8.8M localizations for the sCMOS camera and 6.8M localizations for the CMOS camera. Finally an affine transform was used to convert the CMOS localization positions to the sCMOS coordinate system.

On this test sample there was little to no measurable difference between the performance of the CMOS and the sCMOS camera. This is shown in Fig. 3 where the 2D SMLM images captured by the two cameras of the same field of view appear to be essentially identical in spite of the somewhat worse localization precision of the CMOS camera. All of the microtubules in Fig. 3A are yellow indicating excellent agreement between the sCMOS image (red) and the CMOS image (green). The higher resolution zoomed images in Fig. 3B,C from the two cameras are also essentially identical. Finally, both cameras were able to resolve the hollow structure of the immunostained microtubules in some areas of the image (Fig. 3).

**Figure 3.**
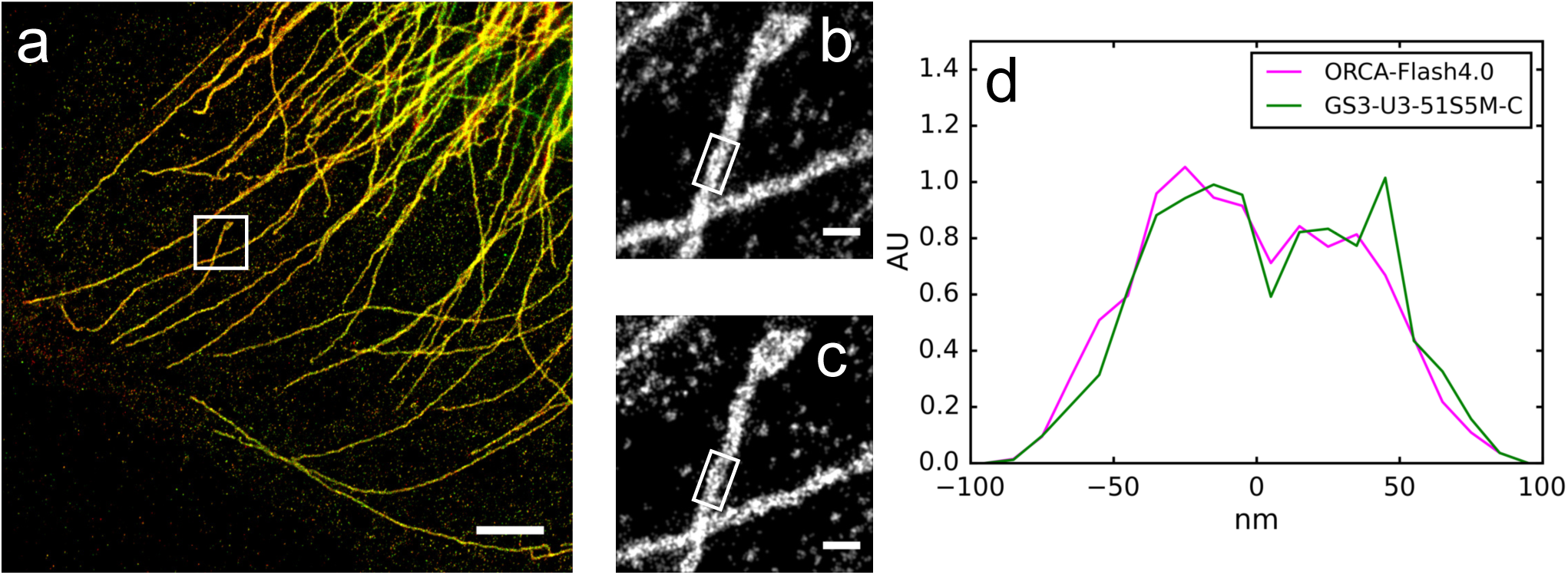
Comparison of 2D SMLM images of Alexa-647 labeled microtubules in U2OS cells acquired with an ORCA-Flash4.0 camera and a GS3-U3-51S5M-C camera. (a) Overlay of the two images (red - ORCA-Flash4.0, green - GS3-U3-51S5M-C). (b) Zoom in of the area white boxed area in (a), ORCA-Flash4.0 camera. (c) Zoom in of the white boxed area in (a), GS3-U3-51S5M-C camera. (d) Histograms of the white boxed areas in (b,c) for the two cameras. Scale bars are 2 μm in (a), 200 nm in (b,c).

Having established that this CMOS camera works for single imaging plane SMLM we next used it for biplane 3D SMLM microscopy^15^. This is a more stringent test of camera performance as in this geometry the fluorescence from a single dye is split by a 50/50 dichroic beam splitter, with half of the light going to each CMOS camera. Lenses L2 in Fig. 1) were adjusted so that the focal planes of the two cameras were offset by ~600 nm. An affine transform was used to map XY positions from the second camera to the first camera. The z dependence of the PSF shape for both cameras was measured by scanning 0.1 μm beads fixed to a coverslip through the focal planes of the two cameras using the piezo objective positioner. Image stacks of the average PSF as a function of z were created only for beads that were visible on both cameras, and that were at least 24 pixels from neighboring beads. A cubic spline was fit to the average PSF image stacks in order to model the shape of the PSF as a function of z for each camera^16^. The cubic splines were used for analysis of biplane SMLM movies with the SMLM cubic spline fitting algorithm described in^16^ adapted for multiplane sCMOS analysis and available here^12^. This adapted algorithm also follows the approach described in^17^ for foreground segmentation instead of using an arbitrary fixed threshold as in the original algorithm.

We were able to acquire biplane 3D SMLM images with this CMOS camera (Fig. 4). The sample again was microtubules in fixed U-2 OS cells labeled with the Alexa-647 flourescent dye. The Alexa-647 dye is one of the brightest SMLM compatible dyes and is expected to emit ~6000 photons on average per switching cycle in our imaging conditions^18^. It may be more difficult to acquire SMLM images of dimmer photo-switchable proteins using this CMOS camera. We also note that we could no longer resolve the hollow structure of the immunostained microtubules, and that our z resolution was relatively low with a FWHM of ~200nm (Fig. 4C). This image was created from 130k frames of data taken at 50 Hz in which SMLM cubic spline fitting analysis identified 24M unique localizations.

**Figure 4.**
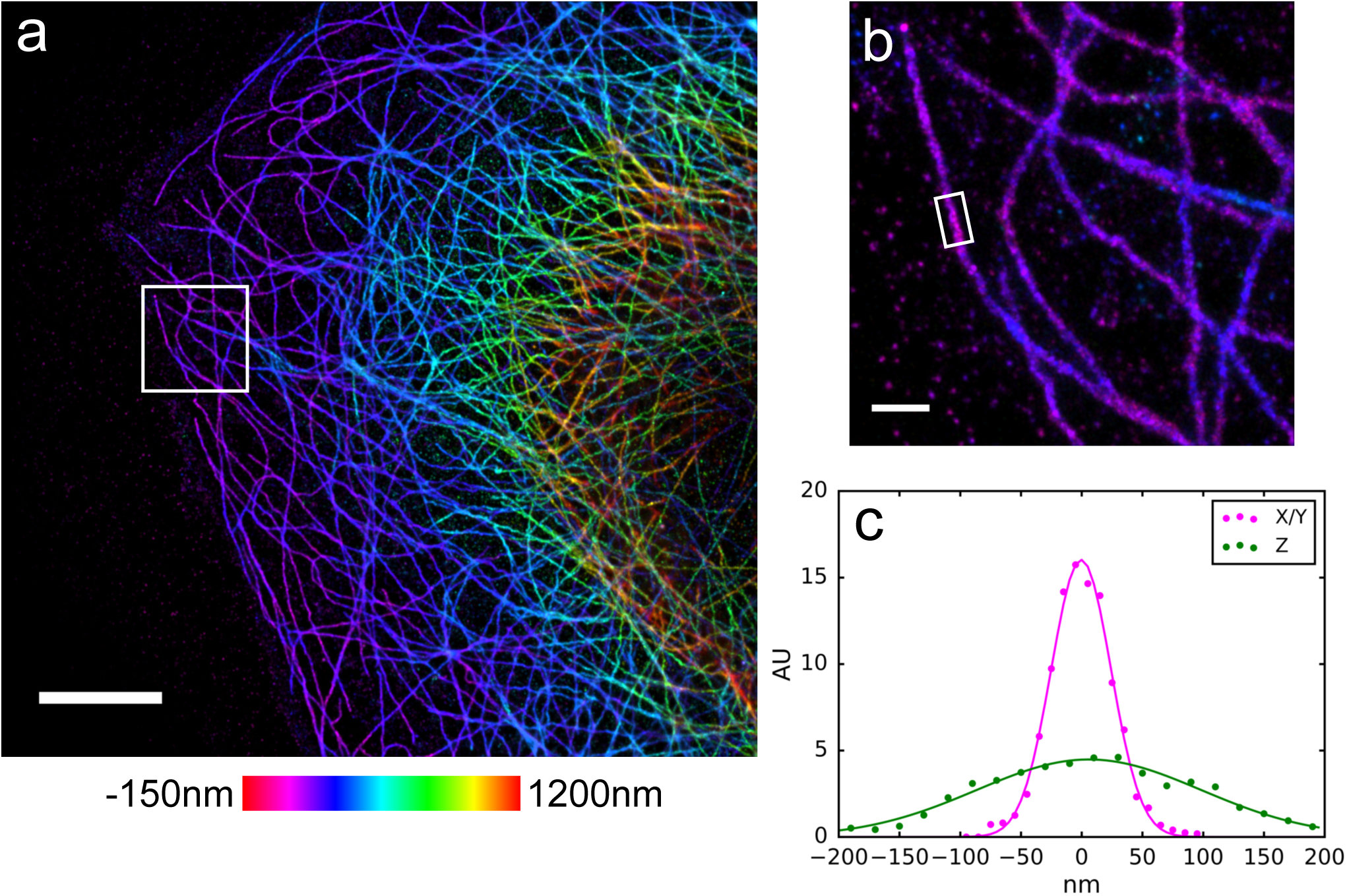
3D SMLM image of Alexa-647 labeled microtubules in U2OS cells taken using two GS3-U3-51S5M-C cameras in a biplane configuration. (a) 3D SMLM image with Z color scale as shown in the color bar. (b) Zoom in of the area white boxed area in (a) with the same Z color scale. (c) Histograms in X/Y and Z of the localizations in the microtubule in the white boxed area in (b). Solid lines are Gaussian fits with σ = 24.5 nm in X/Y and 92 nm in Z. Scale bars are 5 μm in (a), 500 nm in (b).

Next we demonstrate the use of this CMOS camera for quadplane 3D SMLM microscopy. In the quadplane geometry the fluorescence from a single dye is split by 3 50/50 dichroic beam splitters onto 4 different cameras^19^, with each camera receiving 1/4 of the total signal. The imaging planes of the cameras were configured to be separated by ~500 nm enabling the simultaneous acquisition of an ~2.5 μm thick image slice. We followed the same procedure as for biplane imaging, an affine transform was used to map XY positions from the second, third and fourth cameras to the first camera. Cubic splines modeling the PSF for each camera were constructed from z scan image stacks of 0.1 μm beads. The cubic splines were used by the multiplane sCMOS cubic spline algorithm to localize single flourescent dye molecules^12,16^. A quadplane SMLM image of microtubules labeled with the Alexa-647 fluorescent dye is shown in Fig. 5. In this image microtubules that were separated by a distance of ~100 nm in XY were resolved across the entire ~2.5 μm z range. The image was created from 88k frames of data taken at 50Hz in which 8.3M unique localizations were identified.

**Figure 5.**
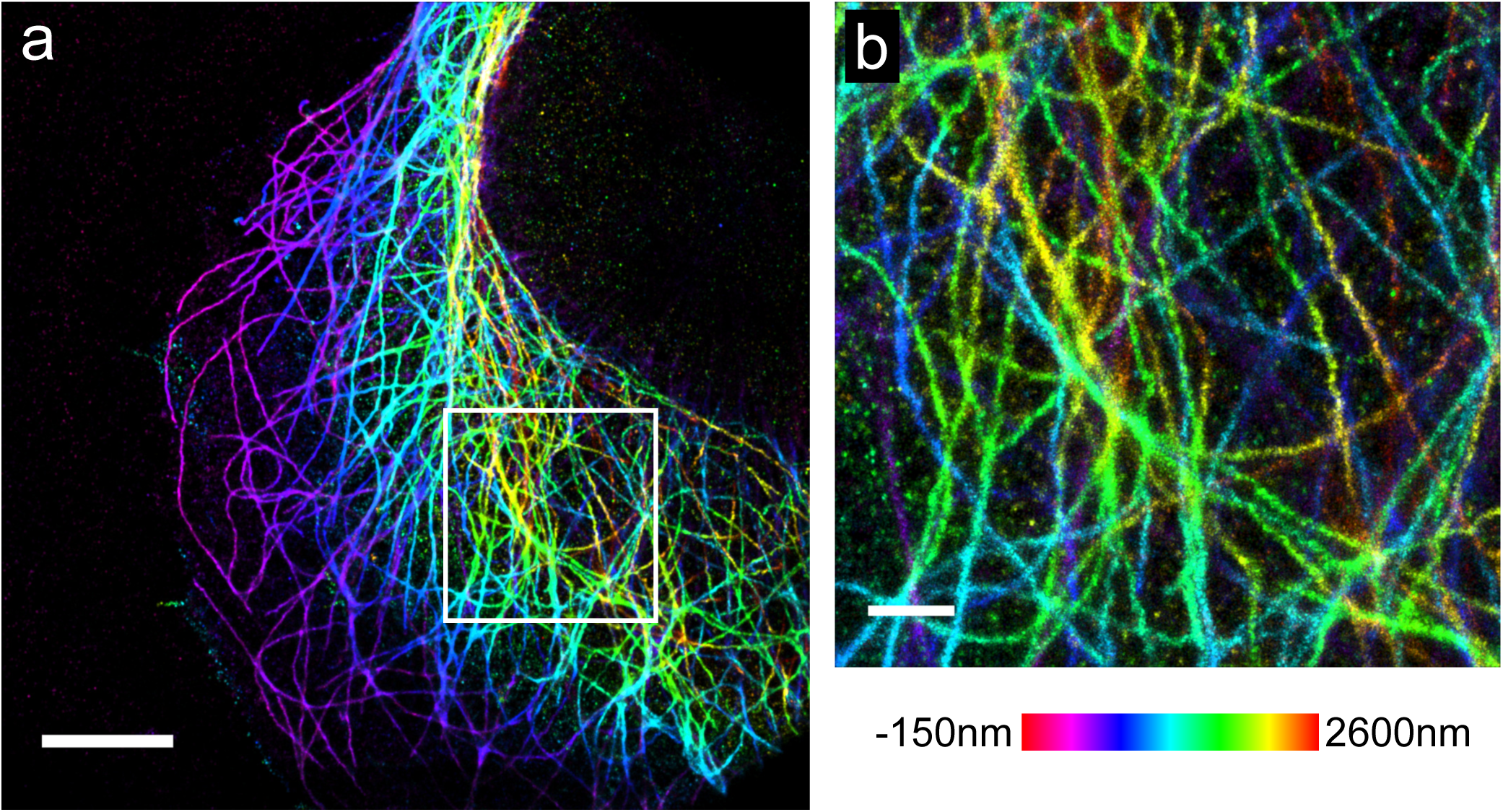
3D SMLM image of Alexa-647 labeled microtubules in U2OS cells taken using four GS3-U3-51S5M-C cameras in a quadplane configuration. (a) 3D SMLM image with Z color scale as shown in the color bar. (b) Zoom in of the area white boxed area in (a) with the same Z color scale. Scale bars are 5 μm in (a), 1 μm in (b).

Finally we demonstrate the use of this CMOS camera for a variant of SR-STORM^10^. Instead of using a prism to spectrally resolve different dye molecules a series of long-pass dichroics was arranged to create 4 different color channels with one camera per color channel. An advantage of this approach is that it tolerates moderately higher localization densities as the dye spectrum is not spread across as many camera pixels. A disadvantage is that the ability to precisely measure the spectrum of a localization is lost. Localization color is instead identified by the relative signal received in each color channel. With this approach it is also possible to simultaneously determine the z position of the localization. The 4 cameras each had relatively small ~100 nm offsets between their focal planes which allowed us to determine z using the same approach as for quadplane imaging. The total z range in this geometry is however limited to about 500 nm as localizations need to be roughly in focus on all 4 cameras simultaneously in order to accurately measure the relative signal in each color channel.

We first imaged dye labeled secondary antibodies non-specifically bound to coverslips. The dyes we choose were DyLight-633, Alexa-647, CF660C and CF680, all of which work well for SMLM imaging as they are bright in their on state, have a duty cycle of ~1:1000 and switch multiple times^10,18^. To measure our ability to discriminate between these dyes, each dye was imaged as a separate sample, then localizations of known dye type were merged into a single data set. We found that by simply computing the first moment of the signal as a function of color channel we could distinguish the dyes with a maximum cross-talk of 5% (DyLight-633 and Alexa-647)(Fig. S1). The maximum cross-talk could be reduced to 2% using a clustering approach. A 4 component vector was created for each localization from the relative signal in each color channel. Then k-means clustering was used to partition the localizations into clusters based on their 4-vectors. Finally, those localizations whose 4-vectors were in the top 20% in terms of distance from the nearest cluster mean were discarded.

A SR-STORM image of microtubules and mitochondria is shown in Fig. 6. In this image microtubules were labeled with the Alexa-647 fluorescent dye and the mitochondrial protein TOMM20 was labeled with the Biotium CF680 dye. The localizations were categorized as Alexa-647 or CF680 based on the distance between their 4-vector and cluster mean vectors. The cluster mean vectors were determined by k-means clustering of an Alexa-674 and a CF680 data set acquired as described in the previous paragraph. We discarded those localizations whose 4-vectors were in the top 20% in terms of distance from the nearest cluster mean.

**Figure 6.**
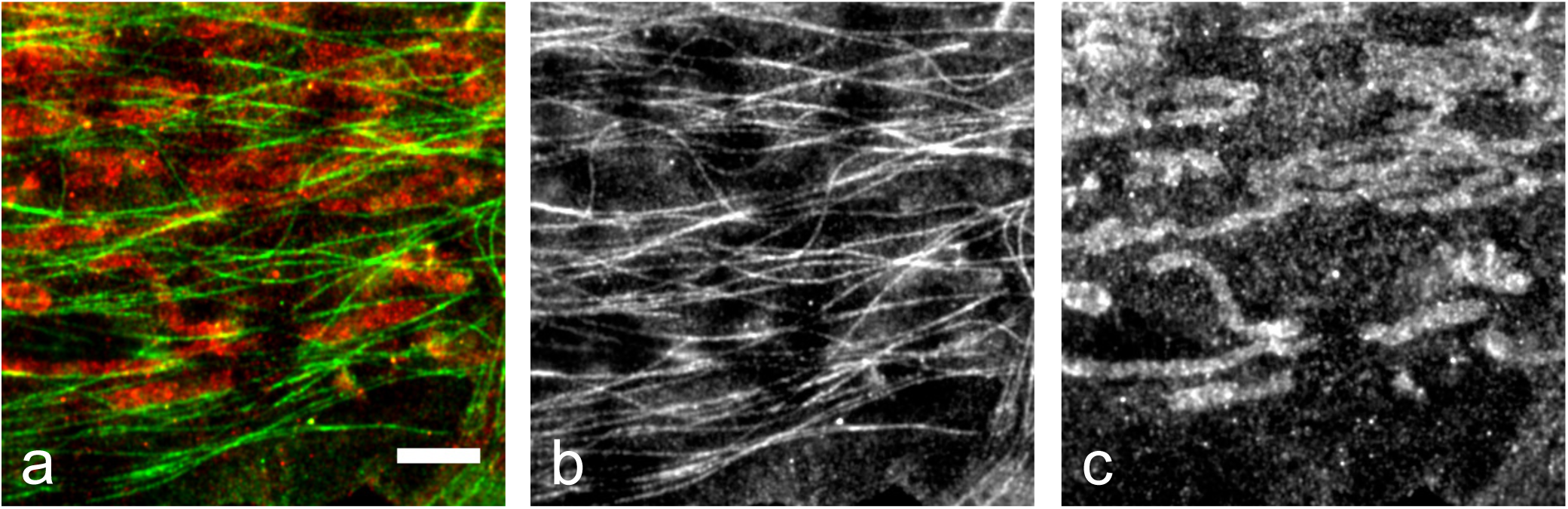
SR-STORM SMLM image of Alexa-647 labeled microtubules and Biotium CF680 labeled mitochondria in U2OS cells. The data was acquired using four GS3-U3-51S5M-C cameras, each detecting a different color channel. (a) Two color image with microtubules shown in green and mitochondria shown in red. (b) Gray-scale image showing the localizations assigned to the Alexa-647 category (microtubule). (c) Gray-scale image showing the localizations assigned to the CF680 category (mitochondria). The scale bar is 2 μm.

## Discussion

In this work we evaluated the use of high end industrial CMOS cameras for SMLM microscopy. We found that they are well suited for the SMLM applications that we tested. However they are not yet as sensitive as the best sCMOS cameras so they may not be a good choice for experiments involving extremely dim dyes. The more than an order of magnitude reduction in camera cost makes it reasonable to consider building microscopy setups that have 4, 8 or possibly even more cameras. This could enable extremely high throughput imaging as well as novel imaging approaches. For example, an octa-plane setup with a ~750 nm spacing between each camera could be used to acquire a z-stack image of an entire cell or thin tissue section in a single exposure. Octa-plane SMLM imaging could be possible with very bright dyes^20^ or approaches like DNA-PAINT^21^. These cameras could also be used for multiplane structured illumination microscopy^22^ or standard epi-fluorescence microscopy in combination with the algorithms to remove sCMOS noise from the final images^23^.

## Methods

### Acquisition Software and Hardware

The setup was controlled with custom software written in Python3 and that used the PyQt5 GUI library. This software was designed to control multiple cameras at once even if they have different chip sizes and/or are from different manufacturers. It supports CMOS cameras from Andor, Hamamatsu and FLIR Imaging. The number and type of cameras that are controlled as well as additional setup functionality is specified in a single XML file in order to make it simpler to use the software to control different setups. This software is open-source and is available on Github^24^.

The four CMOS cameras were controlled using the USB3 interface and a dedicated USB3 PCI Express card (PEXUSB3S7, Startech). For the 512 × 512 AOI that was typically used in the experiments the maximum acquisition rate was ~200 Hz. SMLM movies were streamed directly onto a 1TB PCIe M.2 SSD (MZ-V6E1T0BW, Samsung). In our hands a 7200 RPM hard drive (WD2003FZEX, Western Digital) was not capable of handling the task of streaming data from 4 cameras simultaneously acquiring 512 x 512 pixels images at 50Hz.

### Immunofluorescence and Imaging

Microtubules in fixed U-2 OS cells were fluorescently labeled using the following protocol. Cells cultured in 8 well chambers with a #1.5 coverslip bottom were washed with phospate buffered saline (PBS), permeabilized for 1 minute with a buffer containing 0.1M Pipes, 0.2% triton X-100, 1mM EGTA and 1mM MgCl2, then fixed with a solution of 3% paraformaldehyde and 0.1% glutaraldehyde in PBS (PFA/GA) for 10 minutes. After fixation the cells were washed 3× with PBS then blocked for 15 minutes with 3% BSA, 0.1% triton X-100 in PBS (BB). The anti-beta tubulin primary antibody (ab6046, Abcam) was diluted to 2 μg/ul in BB and incubated for 30 minutes before removal by washing 3× with PBS. The goat anti-rabbit Alexa-647 labeled secondary antibody (A21245, Invitrogen) was diluted to 2 μg/ul in BB and incubated for 30 minutes before removal by washing 3× with PBS. The sample was then post-fixed for 10 minutes with PFA/GA, washed 3× with PBS and stored dry until use.

Microtubules and mitochondria in fixed U-2 OS cells were fluorescently labeled using the same protocol as for microtubule labeling with the following modifications. The initial permeabilization step was removed, after the PBS wash the cells were immediately fixed with PFA/GA. Primary antibodies were rat anti-tubulin (ab6160, Abcam) and rabbit anti-TOMM20 (ab78547, Abcam) both at 2 μg/ul in BB. Secondary antibofies were Alexa-647 goat anti-rat (A21247, Life technologies) and CF680 donkey anti-rabbit (20820-50ul, Biotium) at 2 μg/ul in BB.

Imaging was performed in an oxygen scavenging buffer consisting of 100mM Tris (pH 8.0), 50mM NaCl, 0.5 mg/ml^−1^ glucose oxidase, 40μg/ml^−1^ catalase, 10% glucose and 143mM BME.

## Acknowledgements

This work was supported by the Center for Advanced Imaging at Harvard University. We thank the Zhuang lab for assistance with cell culture, the use of wet bench space and the use of some reagents. Some of the computations in this paper were run on the Odyssey cluster supported by the FAS Division of Science, Research Computing Group at Harvard University.

## Author contributions statement

H.B. conceived the experiments, conducted the experiments, analysed the results and wrote the manuscript.

## Additional information

**Supplementary information accompanies** this paper at; **Competing financial interests** The authors declare that they have no competing interests.

---

**Figure S1.**
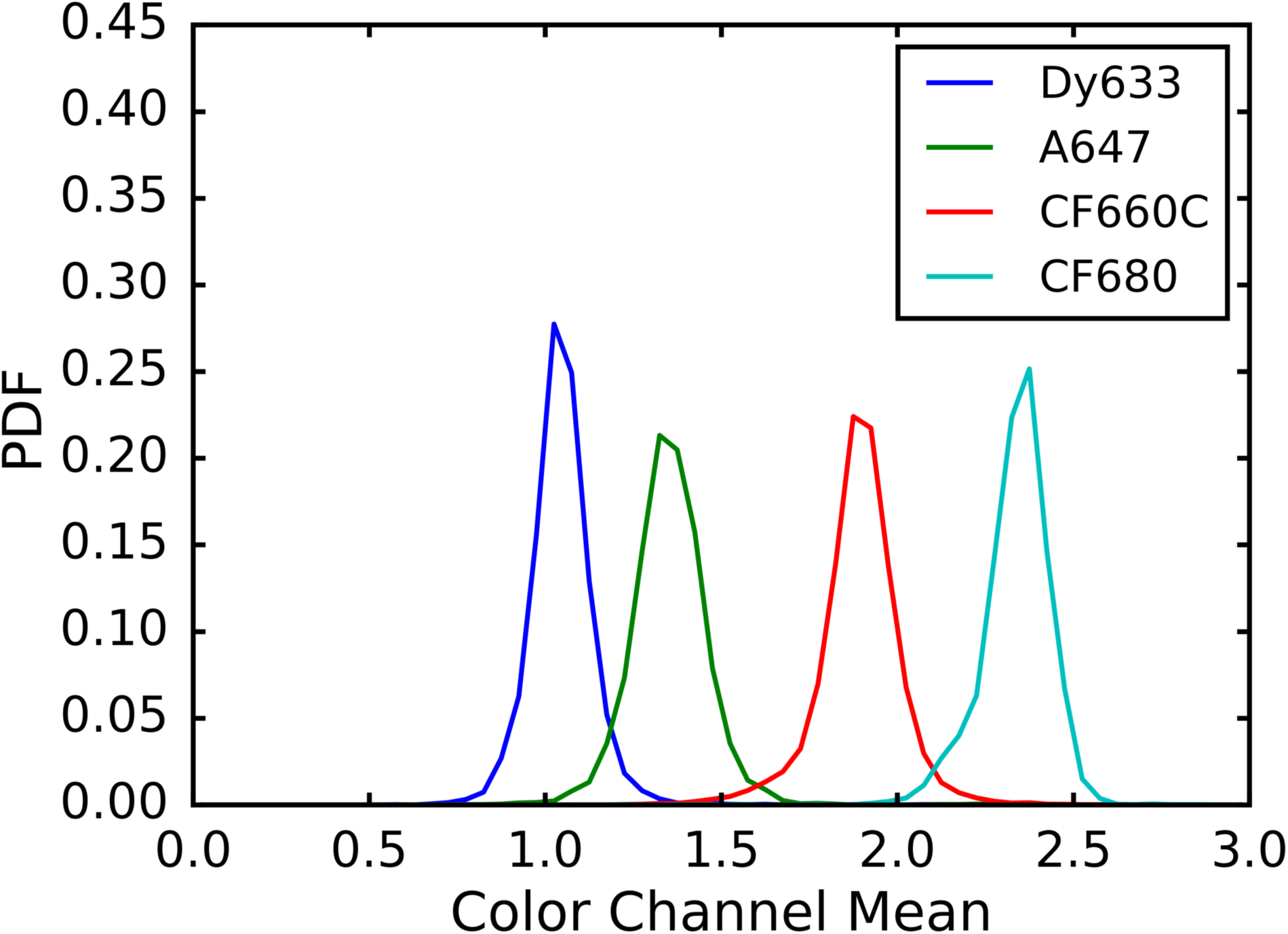
Probability distribution function of the dye color channel mean. Dye labeled secondary antibodies were diluted in PBS and non-specifically bound to coverslips. A single 500 frame STORM movie was taken of each type of dye and analyzed with the multi-channel spline fitting software. For each localization the color channel mean statistic was calculated by multiplying the localizations height in channel by the channel number and dividing by the total of the heights. This statistic is the first moment of the dye heights with respect to color channel number. The color channels were organized from shortest to longest wavelength with channels covering approximately the following wavelengths, channel0 647-670 nm, channel1 670-695 nm, channel2 695-720 nm, channel3 720+ nm. Localizations with a total height of less than 200 e-were removed from the analysis. Dye abbreviations are Dy633 for DyLight 633, A647 for Alexa 647, CF660C for Biotium CF660C and CF680 for Biotium CF680.

## References

1. Betzig, E. et al. Imaging intracellular fluorescent proteins at nanometer resolution. Sci. 313, 1642–1645 (2006).

2. Hess, S. T., Girirajan, T. P. K. & Mason, M. D. Ultra-high resolution imaging by fluorescence photoactivation localization microscopy. Biophys. J. 91, 4258–4272 (2006).

3. Rust, M. J., Bates, M. & Zhuang, X. Sub-diffraction-limit imaging by stochastic optical reconstruction microscopy (STORM). Nat. Methods 3, 793–795 (2006).

4. Alamada, P., Culley, S. & Henriques, R. PALM and STORM: Into large fields and high-throughput microscopy with sCMOS detectors. Methods 88, 109–121.

5. Huang, F. et al. Video-rate nanoscopy enabled by sCMOS camera-specific single-molecule localization algorithms. Nat. Methods 10, 653–658 (2013).

6. Legant, W. R. et al. High-density three-dimensional localization microscopy across large volumes. Nat. Methods 13, 359–365 (2016).

7. Sigal, Y. M., Speer, C. M., Babcock, H. P. & Zhuang, X. Mapping synaptic input fields of neurons with super-resolution imaging. Cell 163, 493–505 (2015).

8. Ma, H., Fu, R., Xu, J. & Liu, Y. A simple and cost-effective setup for super-resolution localization microscopy. Sci. Reports 7 (2017).

9. Kwakwa, K. et al. easySTORM: a robust, lower-cost approach to localisation and TIRF microscopy. J. Biophotonics 9, 948–957 (2016).

10. Zhang, Z., Kenny, S. J., Hauser, M., Li, W. & Xu, K. Ultrahigh-throughput single-molecule spectroscopy and spectrally resolved super-resolution microscopy. Nat. Methods 12, 935–938 (2015).

11. Huang, B., Wang, W., Bates, M. & Zhuang, X. Three-dimensional super-resolution imaging by stochastic optical reconstruction microscopy. Sci. 319, 810–813 (2008).

12. storm-analysis. URL https://github.com/ZhuangLab/storm-analysis.

13. Mortensen, K. I., Churchman, L. S., Spudich, J. A. & Flyvbjerg, H. Optimized localization analysis for single-molecule tracking and super-resolution microscopy. Nat. Methods 7, 377–381 (2010).

14. Tokunaga, M., Imamoto, N. & Sakata-Sogawa, K. Highly inclined thin illumination enables clear single-molecule imaging in cells. Nat. Methods 5, 159–161 (2008).

15. Juette, M. F. et al. Three-dimensional sub-100nm resolution fluorescence microscopy of thick samples. Nat. Methods 5, 527–529 (2008).

16. Babcock, H. P. & Zhuang, X. Analyzing single molecule localization microscopy data using cubic splines. Sci. Reports 7 (2017).

17. Tang, Y. et al. SNSMIL, a real-time single molecule identification and localization algorithm for super-resolution fluorescence microscopy. Sci. Reports 5 (2015).

18. Dempsey, G. T., Vaughan, J. C., Chen, K. H., Bates, M. & Zhuang, X. Evaluation of fluorophores for optimal performance in localization-based super-resolution imaging. Nat. Methods 8, 1027–1036 (2011).

19. Prabhat, P., Ram, S., Ward, E. S. & Ober, R. J. Simultaneous imaging of several focal planes in fluorescence microscopy for the study of cellular dynamics in 3D. Proc. SPIE 6090, 115–121 (2006).

20. Vaughan, J. C., Jia, S. & Zhuang, X. Ultrabright photoactivatable fluorophores created by reductive caging. Nat. Methods 9, 1181–1184 (2012).

21. Jungmann, R. et al. Multiplexed 3D cellular super-resolution imaging with DNA-PAINT and exchange-PAINT. Nat. Methods 11, 313–318 (2014).

22. Abrahamsson, S. et al. Multifocus structured illumination microscopy for fast volumetric super-resolution imaging. Biomed. Opt. Express 8, 4135–4140 (2017).

23. Liu, S. et al. sCMOS noise-correction algorithm for microscopy images. Nat. Methods 18, 760–761 (2017).

24. storm-control. URL https://github.com/ZhuangLab/storm-control.

